# Can you have multiple attentional templates? Large-scale replications of Van Moorselaar, Theeuwes and Olivers (2014) and Hollingworth and Beck (2016)

**DOI:** 10.1101/474932

**Authors:** Marcella Frătescu, Dirk Van Moorselaar, Sebastiaan Mathôt

## Abstract

Stimuli that resemble the content of visual working memory (VWM) capture attention. However, theories disagree on how many VWM items can bias attention simultaneously. The multiple-state account posits a distinction between template and accessory VWM items, such that only a single template item biases attention. In contrast, homogenous-state accounts posit that all VWM items bias attention. Recently, Van Moorselaar et al. (2014) and Hollingworth and Beck (2016) tested these accounts, but obtained seemingly contradictory results. Van Moorselaar et al. (2014) found that a distractor in a visual-search task captured attention more when it matched the content of VWM (memory-driven capture). Crucially, memory-driven capture disappeared when more than one item was held in VWM, in line with the multiple-state account. In contrast, Hollingworth and Beck (2016) found memory-driven capture even when multiple items were kept in VWM, in line with a homogenous-state account. Considering these mixed results, we replicated both studies with a larger sample, and found that all key results are reliable. It is unclear to what extent these divergent results are due to paradigm differences between the studies. We conclude that is crucial to our understanding of VWM to determine the boundary conditions under which memory-driven capture occurs.

---

Regardless of whether you search for a keychain or a mustard bottle, an internal representation of the target object is stored in visual working memory (VWM; Bundesen, 1990; Bundesen, Habekost, & Kyllingsbæk, 2005; Desimone & Duncan, 1995). Such *attentional templates* play a crucial role in optimizing search by means of top-down influences. Behavioral studies showed this by having participants maintain a color in memory while performing a visual search task. These studies generally find that search is slower when the display contains a colored distractor that matched the remembered color, relative to a color-unrelated distractor (Olivers, Meijer, & Theeuwes, 2006; Soto & Humphreys, 2008; Soto & Humphreys, 2007). This *memory-driven attentional capture* suggests that visual working memory and visual attention share content-specific representations (Olivers et al., 2006), and leads to the prediction that items interfere with visual search whenever they match the content of the working memory. Put differently, all yellow bottles in the fridge will capture your attention when you search for the mustard.

Theories differ in the number of items that they postulate can take the role of attentional templates. Olivers and colleagues (2011) proposed a multiple-state account of VWM, which posits two distinct states within VWM: an *active state*, in which at most a single item takes the role of attentional template and consequently biases attention towards task-relevant input; and a *passive state*, in which (multiple) items are stored in VWM but do not interact with visual sensory input. As such, it would be possible to store multiple items in VWM, but only one item would affect visual search. This division within VWM also implies a competition between items when multiple stimuli are relevant for a specific task. In this case, competition might even prevent any item from reaching the state of attentional template, thus abolishing memory-driven attentional capture altogether (cf. van Moorselaar, Theeuwes, & Olivers, 2014).

Dombrowe and colleagues (2011) tested whether people can simultaneously maintain two attentional templates. They conducted an experiment in which participants made a sequence of two eye movements to two colored target objects. In one condition, both targets had the same color, thus requiring only a single attentional template; in another condition, the targets had two different colors, thus requiring two attentional templates. Crucially, Dombrowe and colleagues (2011) found that the second eye movement was delayed by about 250-300ms when the targets had different colors. The authors interpreted this delay as a cost associated with switching attentional templates, and thus as evidence that only one attentional template can be active at a time.

Van Moorselaar et al. (2014) also tested the multiple-state account by having participants maintain a color in memory and simultaneously perform a visual search task, a paradigm partially adopted from Olivers, Meijer and Theeuwes (2006). Participants were first presented with a memory task, where they had to remember one, two, three or four colored items located at one of four possible locations on the screen. Next came a search task that required participants to search for a target among distractors and indicate in which direction the target’s line-segment was tilted. Notably, one of the distractors (if one was present) had a particular color: different or identical to one of the colors presented in the memory task. Van Moorselaar et al. (2014) found that memory-matching distractors capture attention, but, crucially, only when participants kept a single item in memory. The effect of memory-driven capture disappeared at higher memory loads. The results are in line with Olivers and colleagues’ (2011) theoretical proposition of a gateway that allows only one item to become the attentional template. Strikingly, the results even suggest that, at higher memory loads, competition between items prevents any item from becoming an attentional template.

Van Moorselaar et al. (2014) also assessed individual VWM capacity with a change-detection task. The authors did this to test whether the effect of memory-driven attentional capture at higher loads differs among individuals with different working memory capacity. According to the multiple states account, the reduction or abolishment of memory-driven attentional capture at higher loads is due to only one item taking the role of an attentional template, a prediction that does not depend on individual memory capacity. In contrast, *homogenous accounts* posit that memory-driven attentional capture should happen for all items held in VWM, as long as they fall within an individual’s memory capacity. In this opposing view, capture at higher loads should decrease gradually, not because of competing attentional templates, but due to the memory representations becoming less precise as their number in VWM increases (Bays, Catalao, & Husain, 2009; Zhang & Luck, 2008).

Beck, Hollingworth and Luck (2012) argue in favor of the homogenous account on the basis of previous studies that showed distributed activity of memory representations within the visual sensory cortex (Harrison & Tong, 2009; Serences, Ester, Vogel, & Awh, 2009). This view posits an advantage of holding one attentional template compared to two in terms of faster visual search; still, Beck and colleagues (2012) think it possible for two (or more) templates to guide attention simultaneously, even though this would come at the cost of a reduced quality of (each of) the maintained items (Bays & Husain, 2008). This then results in a decrease in performance during visual search tasks (Kristjánsson & Kristjánsson, 2018).

Recently, Hollingworth and Beck (2016) tested the homogenous-state account by means of a paradigm that combined a memory task with a visual-search task, similar (but not identical) to the paradigm used by Van Moorselaar et al. (2014). Participants had to keep either one or two colors in VWM, after which a visual-search task ensued; participants had to indicate the orientation of a target among seven other distractors. The type of distractors was manipulated, with the array containing either only uncolored distractors; two distractors both having colors unrelated to items held in VWM; two distractors, one of which had the same color as (one of the) items held in VWM; or two distractors both matching the color of the items held in VWM. Lastly, participants were tested on one of the items in the memory task. Crucially, Hollingworth and Beck (2016) found memory-driven capture, even when participants held two stimuli in working memory, which they interpreted as evidence that multiple items in VWM can bias attention. The presence of memory-driven capture at load 2 contrasts with the results of Van Moorselaar et al. (2014). It also contrasts with a version of the multiple-state account in which competition between VWM items prevents any item from becoming an attentional template at higher memory loads.

Here we present a replication of two studies: Van Moorselaar, Theeuwes and Olivers’ (2014) experiment one and Hollingworth and Beck’s (2016) experiment (gap-location task). Because previous studies on memory-driven capture have produced mixed findings, and were generally based on small samples, we feel that it is important to establish whether memory-driven capture can indeed occur with a memory load of two items (as claimed by Hollingworth & Beck, 2016) or only with a memory load of one item (as claimed by Van Moorselaar et al., 2014). We replicated Van Moorselaar and colleagues’ (2014) first (singleton-shape task) experiment using the original procedure. However, only conditions using memory load 1 and 2 were tested, because the crucial comparison is between these two conditions. To foresee the results, the memory-capture effect was observed at memory load 1, but not at memory load two, as reported by Van Mooreselaar et al. (2014). The replication of Hollingworth and Beck’s (2016) gap-location task used their original procedure, save one modification: we only tested a memory load of 2 items. To foresee the results, we replicated the memory-driven-capture effect at load 2 with both one and two distractors matching the color of the VWM items (i.e., partial and full capture).

## Experiment 1

The aim of Experiment 1 was to replicate the main finding of Van Moorselaar et al. (2014, Exp. 1) that memory-driven capture occurs at a memory load of one item, but disappears at a memory load of two items.

### Methods

Experimental scripts, data, and analysis scripts can be found at: https://osf.io/pwhkc/. The experiment was conducted as part of a course. 20 students, who were enrolled in this course, tested a total of 66 participants, who participated voluntarily and did not receive compensation. The experiment was approved by ethics review board of the Department of Psychology of the University of Groningen (#17364-O). Stimuli were generated with OpenSesame 3.1 (Mathôt, Schreij, & Theeuwes, 2012) using the PsychoPy backend (Peirce, 2007). Students downloaded this software on their own computers, recruited participants themselves, and performed testing in any environment that they considered suitable. (That is, the data was collected “in the wild”, and not in a traditional laboratory setting.) Participants provided verbal informed consent prior to participation, and were allowed to (and frequently did) abort the experiment whenever they wanted.

The task, illustrated in Figure 1, was closely modeled after Exp. 1 from Van Moorselaar et al. (2014), and implemented by the original author of that study (DM). Each trial started with a 750 ms black fixation cross. Next, a 1000 ms memory display was shown with one (*Load 1*) or two (*Load 2*) colored squares placed at a random subset of four possible locations centered on the intercardinal axes. All stimuli were presented at a distance of 200 px from the central fixation point, and were about 50 px in diameter. Memory colors were randomly selected from five color categories (red, green, yellow, blue, and purple); each color category had nine different exemplars with different combinations of hue and chroma, but all with a similar brightness (except for yellow, which was overall brighter to make it appear less brown). The memory display was followed by a 1250 ms fixation display, followed in turn by the search display.

**Figure 1.**
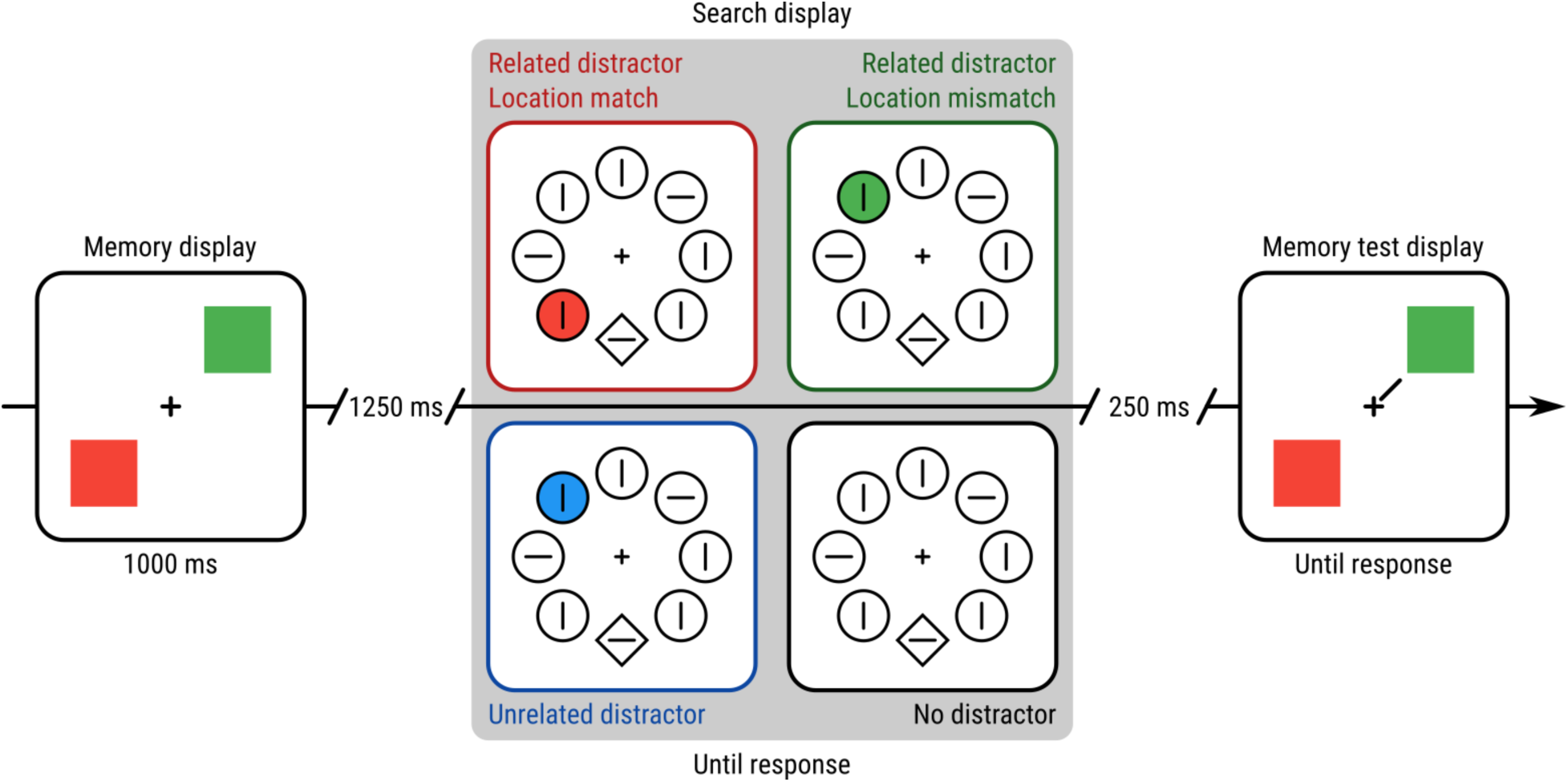
A schematic example trial of Experiment 1. This example demonstrates a Load 2 trial, because the participant needs to remember two colors. For visualization, all Distractor conditions are shown for the search display. However, only one Distractor condition occurred on each trial.

Each search display consisted of one white diamond-shaped target and seven white disk-shaped distractors, all with a leftward or rightward-tilted white line-segment in their center. Stimuli were presented on an imaginary circle around the central fixation dot, also with a 200 px radius, such that the location of the distractors at the intercardinal locations matched the four possible memory locations. The location of the target was randomly selected, with the restriction that it was never displayed on one of the memory locations of that trial. Participants used the arrow buttons on the keyboard to indicate as fast and accurately as possible whether the target line-segment was tilted leftward or rightward from a vertical orientation. In case of an incorrect response, the 250 ms fixation display that followed the search display was replaced by the word “incorrect” displayed in red at the center of the screen.

On 75% of the trials one of the distractors, located at one of the memory locations of that trial, was replaced by a colored disk with a color selected from the memory color pool. There were four distractor conditions, randomly mixed within blocks. In the *Unrelated Distractor* condition, the color category of the distractor was different from the color categories present in the memory display. In the *Color Related / Location Match Distractor* condition, the distractor color matched the memory color that was presented on that location in the preceding memory display. In the *Color Related / Location Mismatch Distractor* condition, the distractor color also matched one of the memory colors, but the colored distractor now appeared at the position of one of the other memory colors. (In case of load 1, this meant that the distractor was placed on an empty memory location.) Finally, in the *No Distractor* condition there was no colored distractor present in the search display.

(Although the colored distractor could be shown at all four possible memory locations, throughout the entire experiment one of these locations had a higher probability (66.7%), and this location was varied between subjects. Trials on which the color distractor appeared at these highly probably locations were called *Suppressed*. The other trials were called *Non Suppressed*. This Suppression factor was included to replicate a study by Wang and Theeuwes (2018) on statistical learning of distractor suppression. The results of this (successful) replication can be found on the Open Science Framework. For the purpose of the current manuscript, we note only that Suppression did not interact with any of the factors of interest here, and do not consider it further.)

The trial sequence ended with a change-detection memory test, in which colored squares were presented on the memory locations of that trial. On half of the trials these squares were identical to the memory display, while on the other half of the trials the color of one of the squares changed to another exemplar from the same color category. A one-pixel thick line cued one square, and participants were instructed to indicate with a key press whether the cued square changed or was identical to the memory item (“C” for change and “N” for change).

All participants completed 15 practice trials and 10 experimental blocks of 48 trials each. Within an experimental block, each Distractor Condition was present six times for each of the memory loads, randomly mixed, resulting in 30 observations per condition. In between blocks, participants were encouraged to take a short break, while they received feedback on reaction times (RTs) and accuracy in the search task, and their accuracy in the memory task.

## Results

### Trial and participant exclusion criteria

Exclusion criteria were based on, but not identical to, Van Moorselaar et al. (2014). Data analysis was performed by SM, based on clarifications by the original author (DM).

First, all trials with a reaction time of less than 200 or more than 5000 ms were excluded. Next, for each participant separately, trials on which the reaction time (RT) deviated more than 2.5 SD from that participant’s mean correct RT were excluded. Next, participants were excluded if: their mean RT deviated more than 2.5 SD from the grand mean RT; or their performance on the memory task was not significantly different from chance (*p* ≥ .05), as determined by a *Χ*^2^ test. (Two participants performed far below chance on the memory task. These were not excluded based on the assumption that they had inadvertently reversed the response rule but had otherwise performed the task correctly.)

56 participants (of 66) and 26,180 trials (of 32,670) remained for further analysis. All results described below are robust to different trimming procedures, including the one used in Van Moorselaar et al. (2014).

#### Effects of Memory Load and Distractor Condition on Search Performance

We first conducted a Bayesian Repeated Measures Analysis of Variance (ANOVA) with per-participant mean RT as dependent variable, and *Distractor Condition* and *Memory Load* as independent variables. We used the Inclusion Bayes Factor based on matched models (“Baws Factor”) to assess the evidence for individual effects (Mathôt, 2017). The labels for strength of evidence are those proposed by Jeffreys (1961), cited by (Wetzels et al., 2011). All tests were conducted in JASP 0.8 with the default settings.

**Figure 2.**
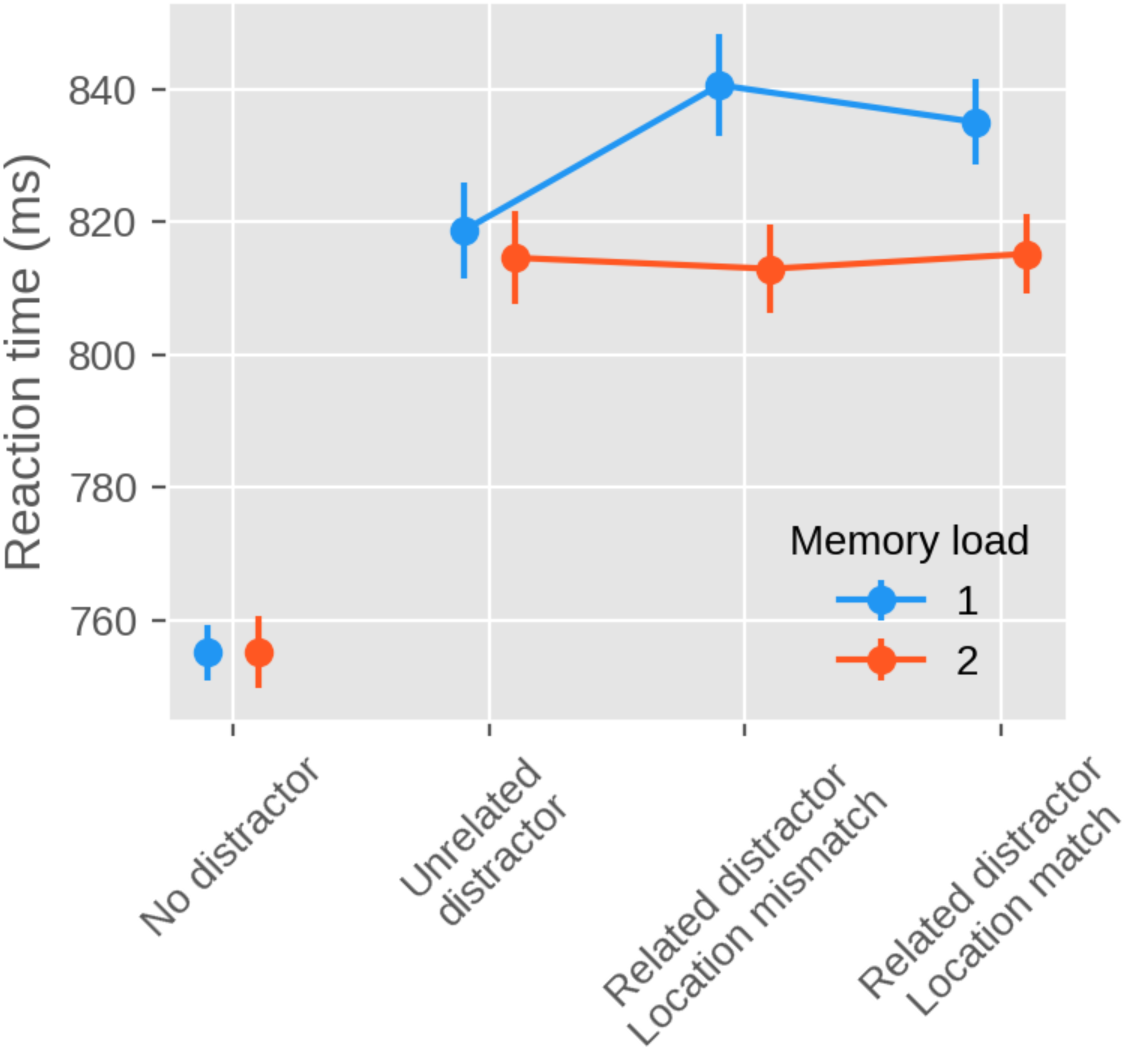
Main results of Experiment 1. Mean response time as a function of Distractor Condition (x axis) and Memory Load (different lines).

There was decisive evidence for an effect of *Distractor Condition* (BF_10_ = 6.00×10^53^), reflecting that participants were overall fastest in the *No Distractor* condition, slower in the *Unrelated Distractor* condition, and slowest of all in the two *Related Distractor* conditions. And there was decisive evidence for an effect of *Memory Load* (BF_10_ = 589.48), reflecting that participants were overall slower on *Load 1* than *Load 2* trials. Crucially, there was substantial evidence for a *Distractor Condition* × *Memory Load interaction* (BF_10_ = 7.99), reflecting that there was a difference between the *Related Distractor* conditions and the *Unrelated Distractor* condition on *Load 1* trials, but not on *Load 2* trials (tested in more detail below).

To further characterize the crucial *Distractor Condition × Memory Load* interaction, which was also reported by Van Moorselaar et al. (2014), we conducted a set of two-tailed Bayesian Paired Samples T-Tests. In the *Load 1* condition, there was strong evidence for higher RTs on *Color Related / Location Match Distractor* trials than on *Unrelated Distractor* trials (BF_10_ = 36.51), and decisive evidence for higher RTs on *Color Related / Location Mismatch Distractor* trials than on *Unrelated Distractor* trials (BF_10_ = 1.19×10^5^). In contrast, in the *Load 2* condition, there was substantial evidence for similar RTs on *Color Related / Location Match Distractor* trials and on *Unrelated Distractor* trials (BF_10_ = 0.163), and substantial evidence for similar RTs on *Color Related / Location Mismatch Distractor* trials and on *Unrelated Distractor* trials (BF_10_ = 0.117). In summary, and consistent with Van Moorselaar et al. (2014), we found memory-driven attentional capture only for a memory load of one item, and not at all for a memory load of two items.

As shown in Figure 3, this pattern was fairly consistent across participants, such that the majority (43 of 56) of the participants showed memory-driven capture in Load 1 condition, and half (28 of 56) of the participants showed memory-driven capture in the Load 2 condition, as would be expected by chance.

**Figure 3.**
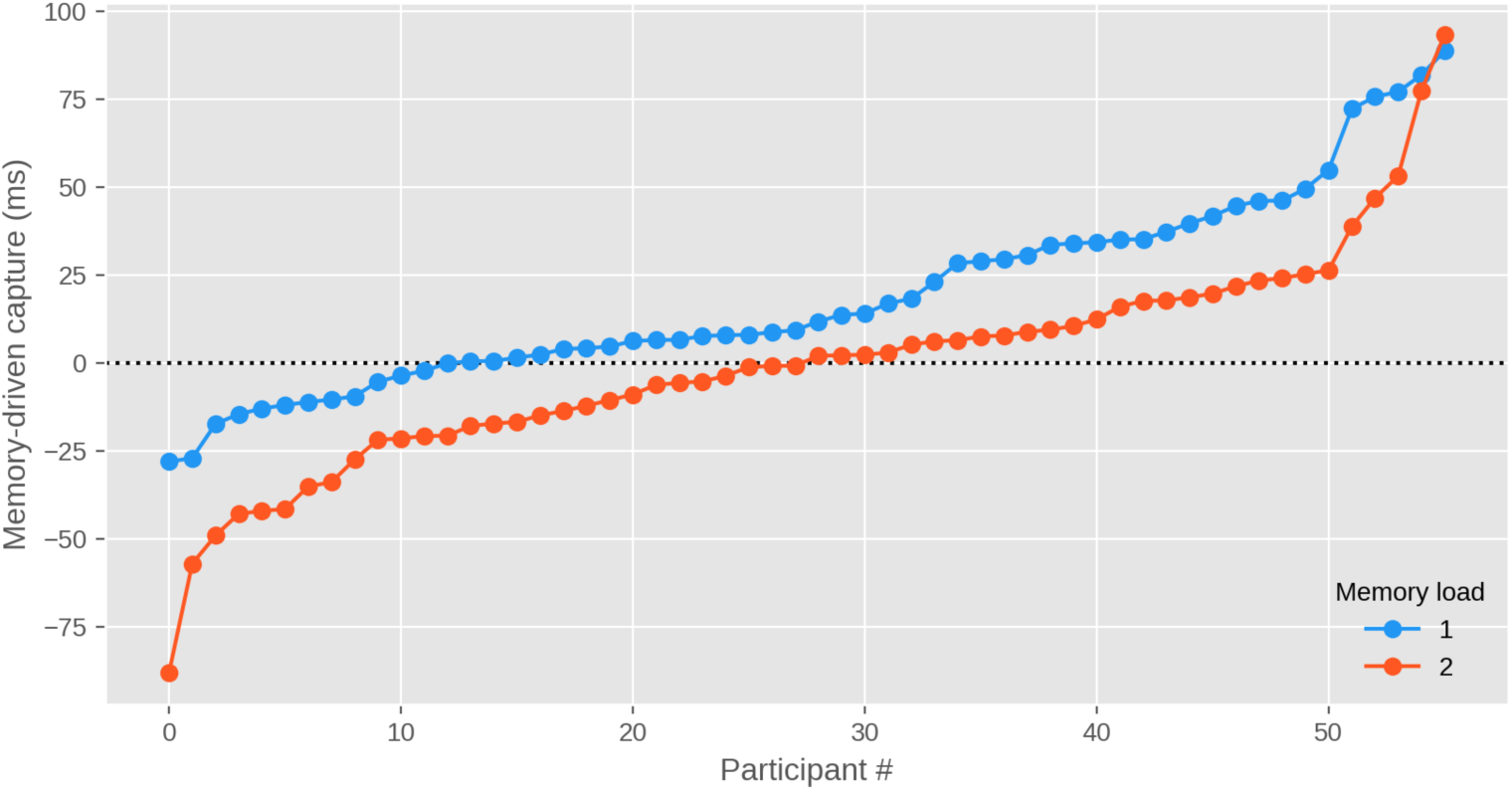
Individual results for Experiment 1. The strength of memory-driven attentional capture for individual participants on Load 1 and Load 2 trials. Memory-driven attentional capture is the difference in response time between the Unrelated Distractor condition and the average of the Color Related / Location Mismatch and Color Related / Location Match conditions. Data points are rank-ordered by effect size, separately for Load 1 and Load 2.

#### Effect of Memory Load on Memory Performance

Finally, as shown by a Bayesian Paired-Samples Two-Tailed T-Test (BF), there was decisive evidence for higher memory accuracy on *Load 1* (76%) than *Load 2* (67%) trials (BF_10_ = 3.52×10^20^).

### Discussion

We successfully replicated the key finding of Van Moorselaar et al. (2014): In their paradigm, memory-driven capture reliably occurs with a memory load of one item, but is completely abolished with a memory load of two items.

## Experiment 2

The aim of Experiment 2 was to replicate the main finding of the gap-location task of Hollingworth and Beck (Hollingworth & Beck, 2016) which showed, in contrast to Van Moorselaar et al. (2014) and our own Experiment 1, that memory-driven capture does occurs at a memory load of two items. A detailed preregistration of this experiment is available at https://osf.io/9k26n/.

The overall approach of Exp. 2 was similar to that of Exp. 1. Only differences are described below.

Students tested a total of 81 participants as part of a course. Stimuli were generated with OpenSesame 3.2 (Mathôt et al., 2012) using the Expyriment backend (Krause & Lindemann, 2014). The task, illustrated in Figure 4, was closely modeled after the gap-location task used by Hollingworth and Beck (2016), based on the original script used in that study, and re-implemented by two authors of the current paper (MF and SM). Each trial started with a 500 ms white fixation dot. Next, a 250 ms memory display appeared, containing two colored squares placed at opposite sides of an imaginary circle around a central fixation dot, with a 128 px radius. The memory display was followed by a 750 ms fixation display, followed in turn by the search display.

**Figure 4:**
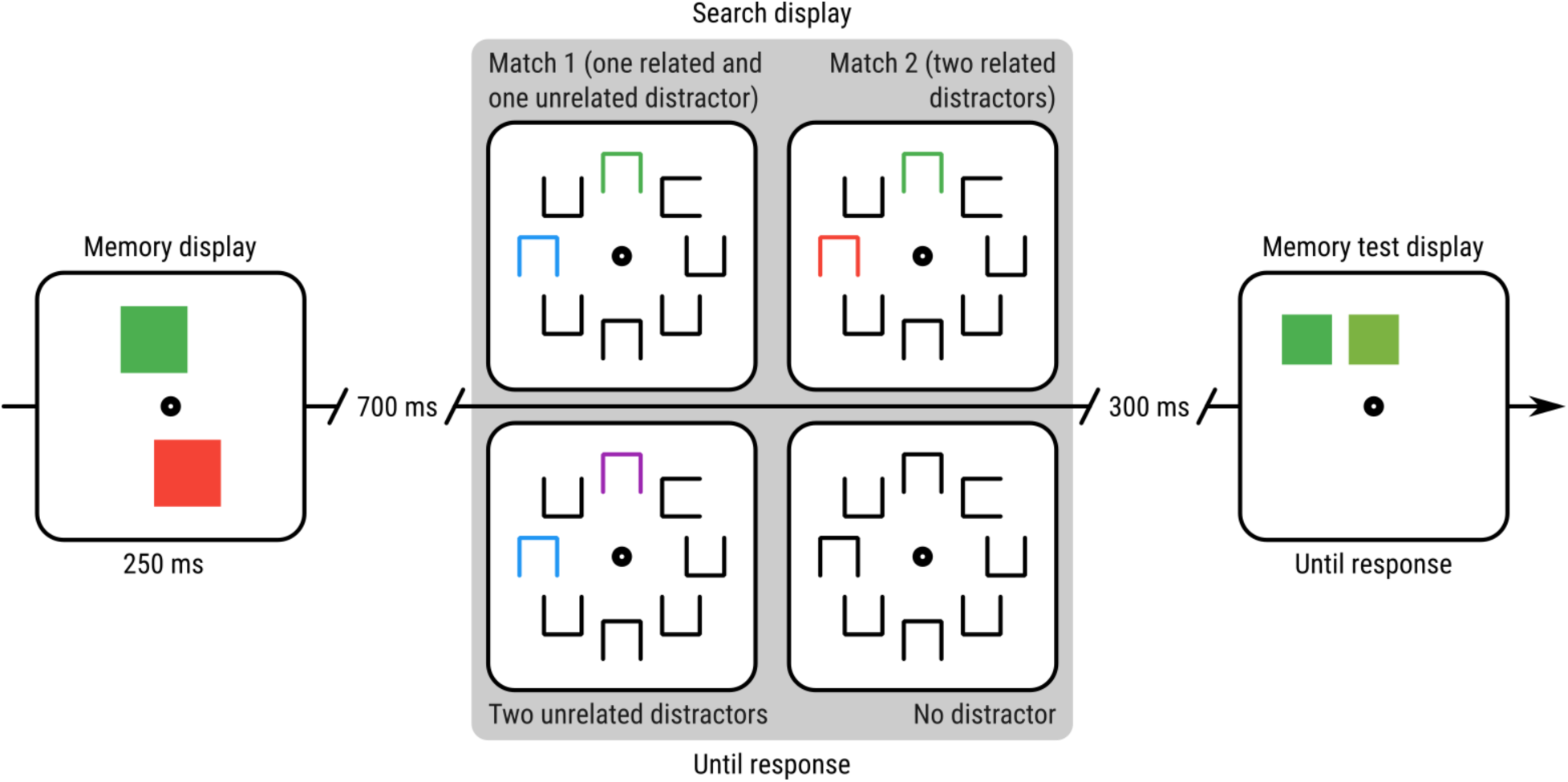
A schematic example trial of Experiment 2. For visualization, all Distractor conditions are shown for the search display.

The search display consisted of eight square-like stimuli, one target and seven distractors, that were open on one side. The target was open on the left or right side. The distractors were open on the upper or lower side. Stimuli were shown on an imaginary circle around the central fixation dot, with a radius of 200 px. Participants used the arrow buttons on the keyboard to indicate as fast and accurately as possible whether the target was open on the left or the right side.

There were four distractor conditions, randomly mixed within blocks, but occurring with different frequencies. In the *Color Unrelated* (20% of trials) condition, two of the distractors were colored (i.e. not white), but their color category differed from the color categories present in the memory display. In the *Match-1* (20% of trials) condition, two colored distractors were shown, one of which matched one of the color categories of the memory display, and one of which did not. In the *Match-2* (20% of trials) condition, two colored distractors were shown, each matching one of the color categories of the memory display. The color match could be exact (e.g. the same shade of green) or categorical (e.g. a slightly different shade of green). Finally, in the *No Distractor* (41% of trials) condition, there were no colored distractors present in the search display.

The trial sequence ended with a color-selection memory test, in which two colored squares were shown side by side. One of the squares exactly matched one of the colors of the memory display; the other square was a categorical match. The squares were shown at the same location as the original color had been shown during the memory display. Participants indicated which of the two squares was an exact match with the memory display by pressing the left or right arrow key. After an incorrect response, the word “incorrect” was shown in red for 300 ms. Finally, after a delay of 300 ms, participants were asked to press the spacebar to start the next trial.

All participants first completed 10 practice trials that contained only the memory task. Next, participants completed 12 practice trials that contained both the memory and search task. Next, they completed 3 experimental blocks of 51 trials each. In between blocks, participants were encouraged to take a short break.

### Results

We used the same exclusion criteria and analyses as in Exp. 1^1^. 56 participants (of 81) and 8,290 trials (of 14,175) remained for further analysis.

We first conducted a Bayesian Repeated Measures ANOVA with per-participant mean RT as dependent variable, and *Distractor Condition* as independent variable (Figure 5). There was decisive evidence for an effect of *Distractor Condition* (BF_10_ = 7.26×10^9^). To characterize this effect, which was also reported by Hollingworth and Beck (2016), we conducted a set of two-tailed Bayesian Paired Samples T-Tests. There was decisive evidence for higher RTs on *Match-2* trials than on *Match-1* trials (BF_10_ = 158.46). There was substantial evidence for higher RTs on *Match-1* trials than *Color Unrelated* trials (BF_10_ = 3.86). Finally, there was strong evidence for higher RTs on *Color Unrelated* trials than on *No Distractor* trials (BF_10_ = 15.98).

**Figure 5.**
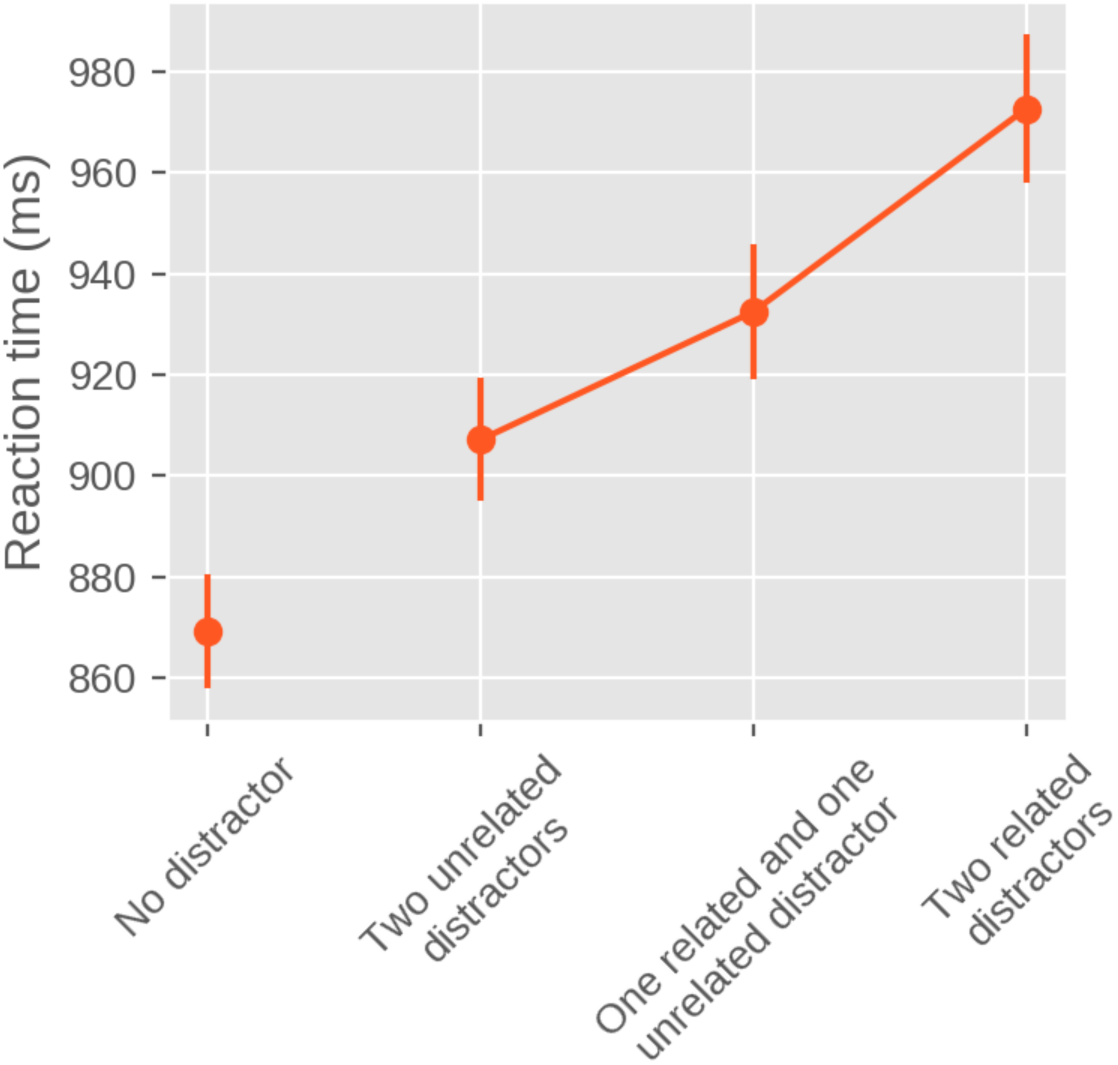
Main results of Experiment 2. Mean response time as a function of Distractor Condition.

As shown in Figure 6, this pattern was fairly consistent across participants, such that the majority of participants (42 of 56) showed stronger memory-driven capture on *Match-2* than *Match-1* trials, and a majority of participants (35 of 56) also showed stronger capture on *Match-1* than *No Distractor* trials.

**Figure 6.**
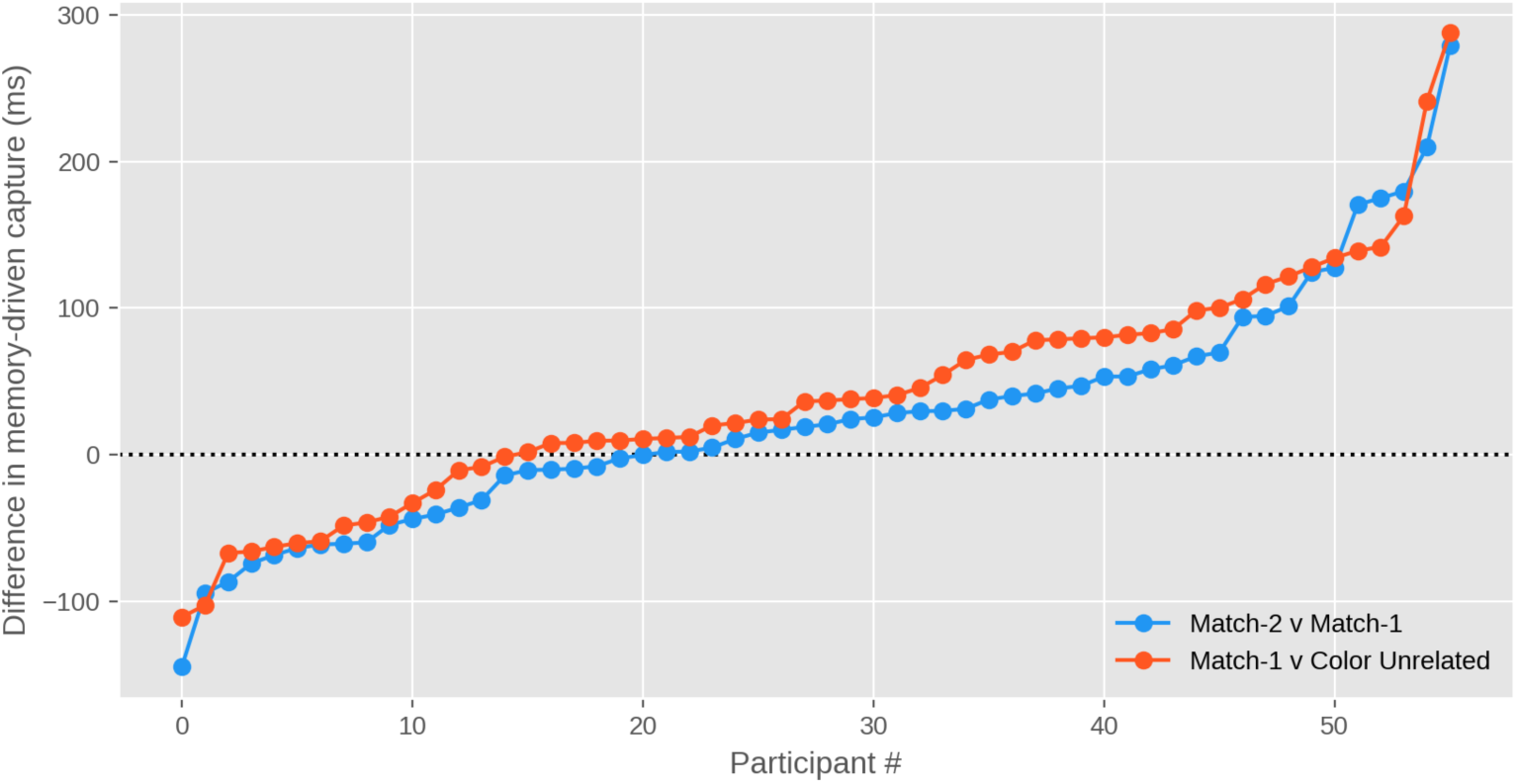
Individual results for Experiment 2. The difference in response time between Match-2 and Match-1 trials (blue) and Match-1 and Color Unrelated trials (red). Data points are rank-ordered by effect size, separately for the two contrasts.

### Discussion

We successfully replicated the key finding of Hollingworth and Beck (2016): In their gap-location task, memory-driven capture reliably occurs with a memory load of two items. And the strength of memory-driven capture is stronger when two distractors match the memorized colors, compared to when only one distractor matches a memorized color.

## General Discussion

The current study successfully replicated two experiments: by Van Moorselaar and colleagues (2014) and by Hollingworth and Beck (2016). First, in replication of Van Moorselaar and colleagues (2014), memory-driven attentional capture was observed when participants kept one item in VWM, an effect well established in the attentional/VWM literature (e.g., D. Soto, Humphreys, & Rotshtein, 2007; David Soto & Humphreys, 2007). As such, a distractor that shared the color of the item held in VWM attracted more attention relative to a distractor that did not share such VWM feature content. Importantly, this effect disappeared at load two; in this case, no memory-driven attentional capture was observed when two items were held in VWM. These results provide support for Olivers and colleagues’ (2011) proposition of a functional distinction in VWM, with an active state in which a memory item biases attention (attentional templates), and an accessory state, where items do not interact with visual search. It is thus possible that when only one item was relevant to the task, it took the role of an attentional template and biased visual search. Conversely, when two equally relevant representations needed to be held in VWM, competition between them prevented either of them from becoming an attentional template, being stored instead as accessory items.

The replication of Hollingworth and Beck’s (2016) experiment showed a strikingly different pattern. There, memory-driven attentional capture was observed for two items held in VWM when one distractor matched the memory content (partial capture) and both distractors matched the memory content (full capture). It thus seems that it was possible for (at least) one attentional template to be instated when two items are held in VWM. The increased time spent on the visual search task when both distractors matched the content of the VWM compared to 1 match, points to the possibility of two attentional templates biasing visual search simultaneously, or a relatively quick switch between equivalently relevant templates. Both cases are incompatible with a version of the multiple state-account that predicts no attentional template role allocated to any of the competing VWM representations. But the simultaneous guidance of two attentional templates is consistent with homogenous accounts predictions of multiple VWM representations biasing attention.

The aforementioned results are puzzling, since both effects are robust, but, at least on the surface, contradictory. Therefore, it is important to mention differences between the two paradigms that could account for discrepancies between results. First, Van Moorselaar and colleagues (2014) had one colored distractor in the visual search task, in both load 1 and 2 conditions of the experiment. On the other hand, Hollingworth and Beck (2016) had two colored distractors. More so, the timing of the two experiments differed; the memory display in Van Moorselaar et al. (2014) lasted for 1000ms (with a subsequent delay of 1250ms), while the memory display in Hollingworth and Beck (2016) lasted for 250ms (with a delay of 700ms). While it is unclear how these paradigm differences could account for the diverging results, they are notable and warrant further investigation. Specifically, future experiments could vary the timing of the memory display (in either of the paradigms) to check whether the effect of memory capture at load 2 still holds – or is abolished as is the case in van Moorselaar and colleagues (2014).

In summary, two different paradigms seemingly provide evidence for two mutually exclusive theories: the multiple-state account that posits that only item can serve as an attentional template at a time, and in its strongest form, that if multiple items are kept in working memory none of them take the role of attentional template; and the homogenous account that posits that multiple items can serve as attentional templates at a time. A single demonstration of at least one attentional template being active with a memory load of two, as provided by Hollingworth and Beck (2016) and also our successful replication, falsifies a strong version of the multiple-state account. On the other hand, a single demonstration of an absence of memory-driven capture with a memory load of two, as provided by Van Moorselaar et al. (2014) and also our successful replication, also suggests that the predictions of the homogenous account do not hold in all situations. This leaves two possibilities: a version of the multiple-state account that allows for at least one attentional template even in situations where multiple items are kept in working memory; or a homogenous account in which all working-memory items can serve as attentional templates, but do not always do so.

## Footnotes

1 Due to a bug in the experimental script, the correctness of the visual-search responses was not logged in Experiment 2. Therefore, analyses are performed across all responses, rather than only correct responses, as we had specified in the pre-registration. To verify that this did not alter the results, we conducted all analyses for Experiment 1 both with and without correct responses, and found that this did not affect any of the results.

